# SpotGraphs: Graph-based analysis of spatially resolved transcriptional data in R

**DOI:** 10.64898/2026.03.12.711347

**Authors:** Alex J. Lee, David E. Sanin

**Author notes:** **Author Contributions Alex J. Lee:** Conceptualization, Methodology, Software, Validation, Formal analysis, Writing - Original Draft, Visualization; **David Sanin:** Conceptualization, Resources, Writing - Review & Editing, Supervision, Funding acquisition.

## Abstract

**Introduction:** Common spatial transcriptomic analysis pipelines in R focus on pre-processing and visualization, while providing limited and indirect methods to leverage true spatially resolved quantification of transcripts. Often, x,y-coordinates in spatial transcriptomics (ST) data are integrated into analysis via “spatially aware” normalization (Salim et al., 2024), clustering methods (Zhao et al., 2021), or the identification of spatially variable genes (Yan et al., 2025). Though useful, these methods do not provide any opportunity for analysts to adjust or interrogate the underlying graphs that define adjacencies between spots in their data. Here, we present SpotGraphs, a package that allows the user a more direct and flexible option to interact with the x,y-coordinates of their ST data in R through the existing igraph infrastructure (Antonov et al., 2023; Csárdi et al., 2025; Csardi & Nepusz, 2006). Similar functionality exists in Python through SquidPy’s graph API (Palla et al., 2022), and we compare results obtained from both packages, demonstrating similar performance. Additionally, we provide a set of tools that are useful for ST data analysis, including a toolkit to filter low quality spots laying on tissue debris, beyond arbitrary thresholds, edit spot-level adjacencies based on spatial clusters, and identify centers or boundaries of user-defined neighborhoods of interest.

## Methods

### Constructing a graph from coordinates

For illustration, we use 10X Visium data collected from oral squamous cell carcinoma samples (Arora et al., 2023). To construct a graph given x,y-coordinates of spatial data, the SpotGraph() function implements two approaches to identify neighboring spots and build an adjacency matrix:

#### 1. Euclidean Distance

We start by calculating a square matrix, M, of Euclidean distances between *n* spots setting a maximum distance (d_max_), to identify which spots are immediately adjacent to each other with the following equation:

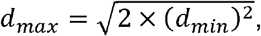

where d_min_ is defined as the minimum Euclidean distance between any two spots. The adjacency matrix defined by M ≤ d_max_ is used to create an igraph object that can be visualized using the original spatial coordinates (Figure 1A -left).

**Figure 1.**
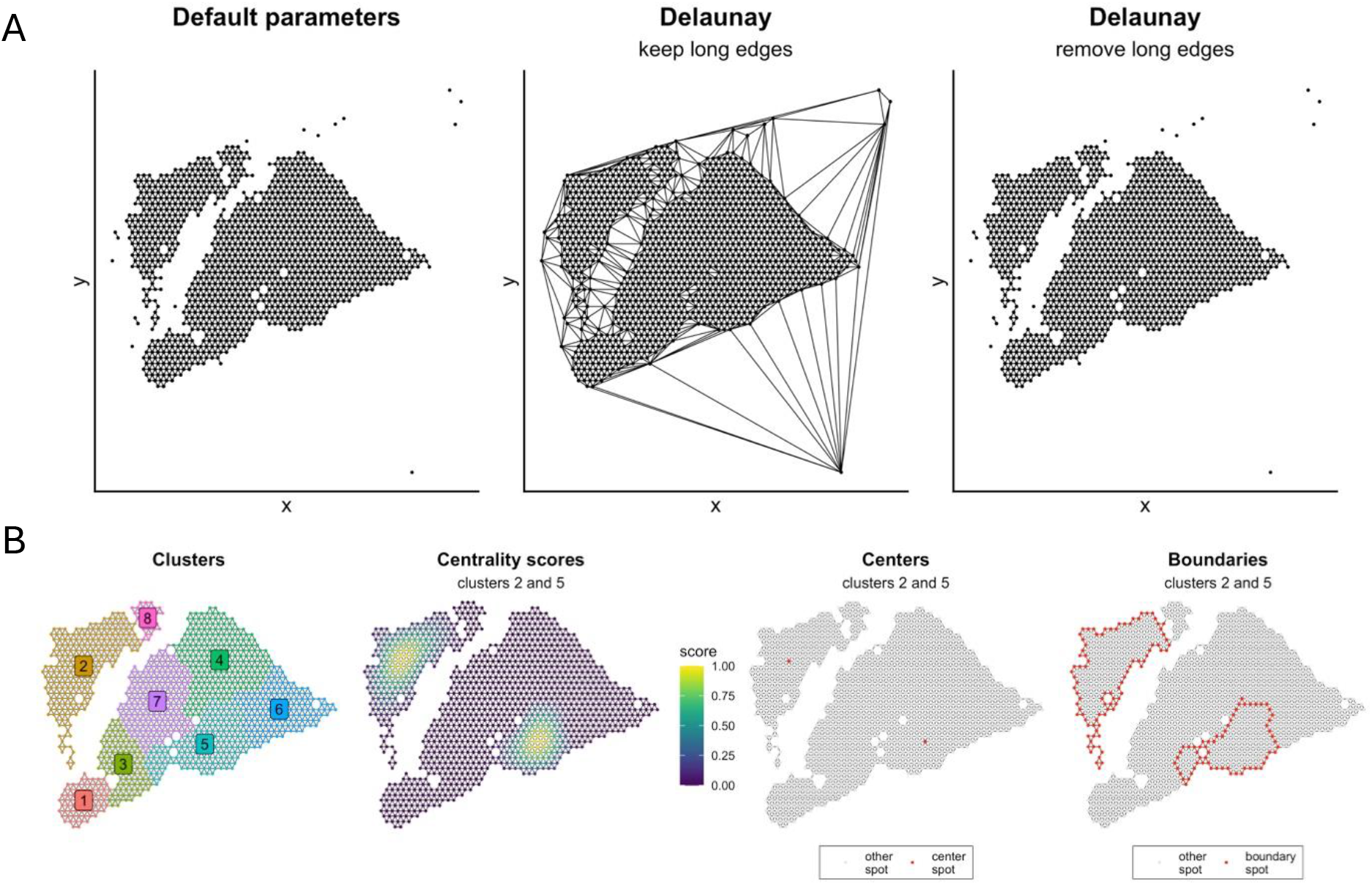
Overview of SpotGraphs functions and methodology. Spot coordinates of 10X Visium data from GSE208253. (A) Results of default network construction based on Euclidean distance, Delaunay triangulation, and Delaunay triangulation followed by removal of edges longer than some user-defined distance threshold. (B) Procedures implemented by the NeighborhoodCenters function. Clusters 2 and 5 are selected to demonstrate centrality scoring, which is used to determine the spot at the center of each spatial cluster. Spots laying at the boundaries of each neighborhood are also identified.

#### 2. Delaunay Triangulation

We use the Delaunay triangulation implementation from the interp R package (Gebhardt et al., 2024). Resulting graphs often contain edges between spots/cells that are not adjacent to each other upon visual inspection (Figure 1A - center). To address this, we include an additional step where edges exceeding some distance, by default d_max_, are removed from the final graph (Figure 1A - right).

The SpotGraph() function returns igraph objects with either strategy, however, the Euclidean distance approach works best when spots are arranged in a grid pattern, either hexagonal (6 adjacent spots) or square (8 adjacent spots). For spots/cells that are not arranged in a grid pattern, we recommend using Delaunay triangulation instead.

### Identifying centers of neighborhoods

A common analysis task in ST analysis is to evaluate the relationship between gene expression patterns and some neighborhood of interest. Two common characteristics that we can use to summarize these neighborhoods are their center point and their boundaries. For example, this may be useful in studying how gene expression varies with distance to the center of each tumor or with distance to some tumor boundary. The NeighborhoodCenters() function makes this type of analysis straightforward by first constructing a network given a set of x,y-coordinates and removing edges that connect spots within the neighborhood and outside of the neighborhood. Eigenvector centrality scores are computed within each neighborhood, and the center point of each neighborhood is defined as the spot(s) with the maximum centrality eigenscore (Figure 1B).

### Availability

All functions, installation instructions, and usage examples can be retrieved from https://github.com/Sanin-Lab/SpotGraphs.

## Benchmarking

To compare the performance of our R-based SpotGraphs package with the existing Python-based SquidPy graph API, we used coordinates from the 13 datasets available through the TENxVisiumData Bioconductor package (Crowell, 2025) to generate graphs with each of 3 methods: grid-based, Delaunay triangulation, and nearest-neighbor (NN), with either SpotGraphs or with SquidPy via the reticulate package (Kalinowski et al., 2025). After summing the number of differences in the upper right triangles of each adjacency matrix, we find that SpotGraphs and SquidPy generate the same adjacency matrix with the grid-based approach, with few differences in the Delaunay triangulation approach, and many differences with nearest-neighbors for edge detection (Figure 2A).

**Figure 2.**
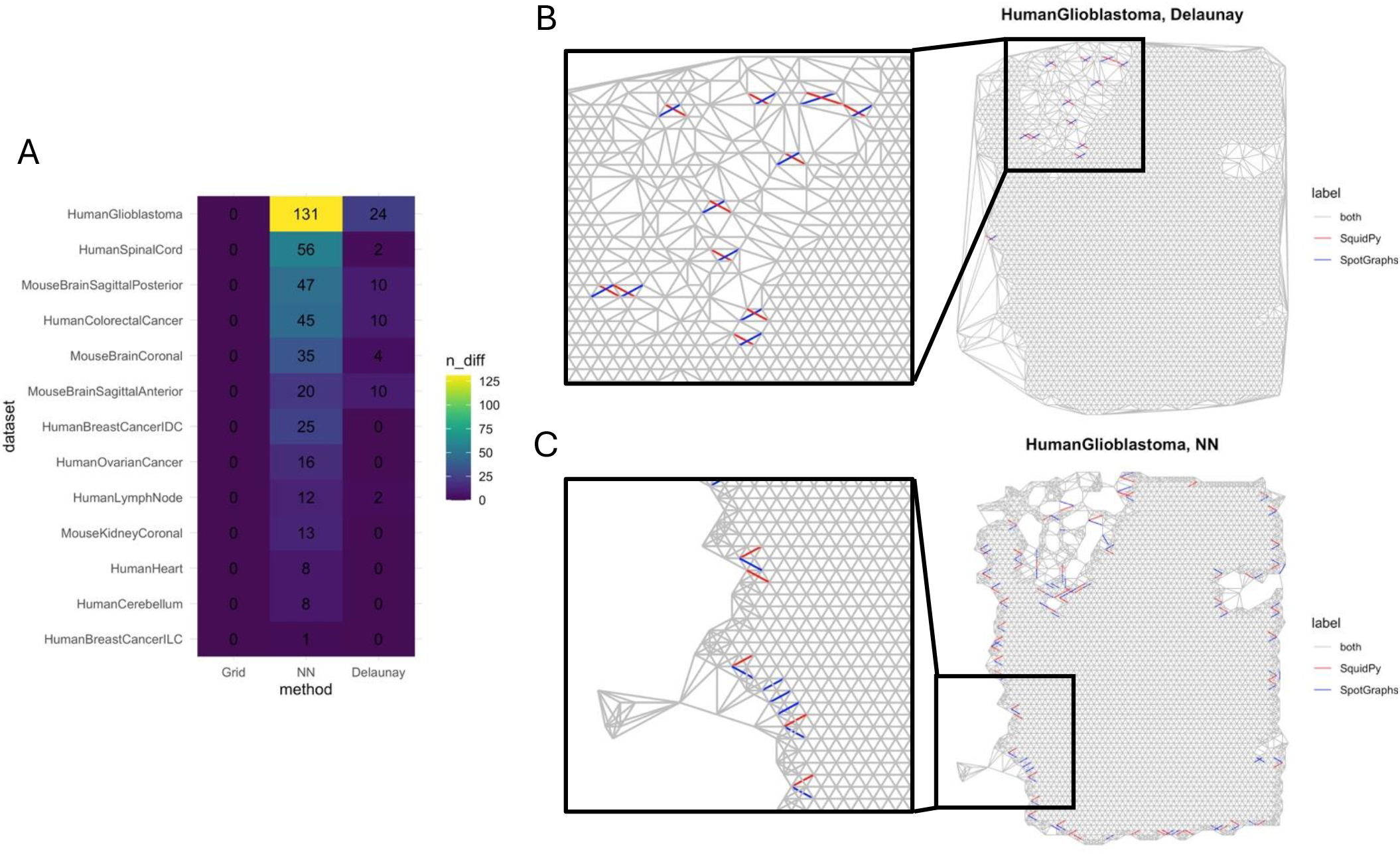
Benchmarking between SpotGraphs and SquidPy’s graph API. (A) The number of differences in the upper right triangles of the adjacency matrices created by either SpotGraphs (for Grid and Delaunay), RANN (for R-based nearest neighbors), or SquidPy. The absence or presence of an edge in one matrix compared to the other matrix is counted in this difference. (B) The spots with corresponding adjacencies plotted for the TENxVisium HumanGlioblastoma dataset. Adjacencies determined by Delaunay triangulation as implemented in either SpotGraphs (via the interp package) or SquidPy. Red indicates edges drawn by SquidPy, but not by SpotGraphs, whereas blue indicates the opposite. (C) The same as in B, but with adjacencies identified by nearest neighbor in RANN or SquidPy with N=6.

Upon further investigation of the HumanGliobastoma dataset, which had the most differences between SquidPy and R-based methods, we find that Delaunay triangulation shows differences where four spots lay on the same circumcircle such that multiple optimal solutions exist that satisfy the properties of Delaunay triangulation (Figure 2B). The nearest-neighbor approach in this dataset shows differences exclusively at the boundaries of the tissue sample (Figure 2C). Despite the differences between the nearest-neighbor implementations, either with SquidPy or RANN (Jefferis et al., 2024), this approach seems generally problematic at tissue boundaries, where it erroneously identifies more neighbors than are actually adjacent to the spots in these regions.

In our benchmarking, we find that SpotGraphs produces identical adjacency matrices to SquidPy with the grid-based approach and comparable adjacency matrices with Delaunay triangulation. We also find that the nearest-neighbor approach tends to overestimate the connections between spots, making it an inappropriate method for determining adjacencies in spatial data.

## Applications

The goal of this package is to provide users access to the igraph infrastructure by providing the x,y-coordinates of their spatial transcriptomics data as input. In general, once an igraph object is created with the SpotGraph() function, various graph-based statistics can be calculated, which can be stored as meta data in the user’s original Seurat object or SpatialExperiment object. We provide a few scenarios where functions made available through SpotGraphs can interact with igraph functions to infer useful statistics about spatial data. Vignettes with relevant code for each scenario are provided in the R package.

### Spot-level filtering

In a situation where count-based spot-filtering has not sufficiently removed spots from uninteresting regions (i.e., areas with no tissue, areas with debris, etc.), the SpotGraph() output can be used to identify poorly connected or isolated spots/cells that should be excluded. Specifically, graphs resulting from this approach can be clustered and visualized using SpatialPlotGraph(), so that users can decide which clusters of spots to analyse further. For example, In the MouseKidneyCoronal sample, we have spots that lay on a fragmented section of the tissue sample that are likely low quality, but are indistinguishable by typical quality check metrics such as read count (Figure 3A - left image). Following graph creation and clustering (Figure 3A - center), we identify and remove spots with a low degree (i.e., few adjacent spots) before proceeding with downstream analyses.

**Figure 3.**
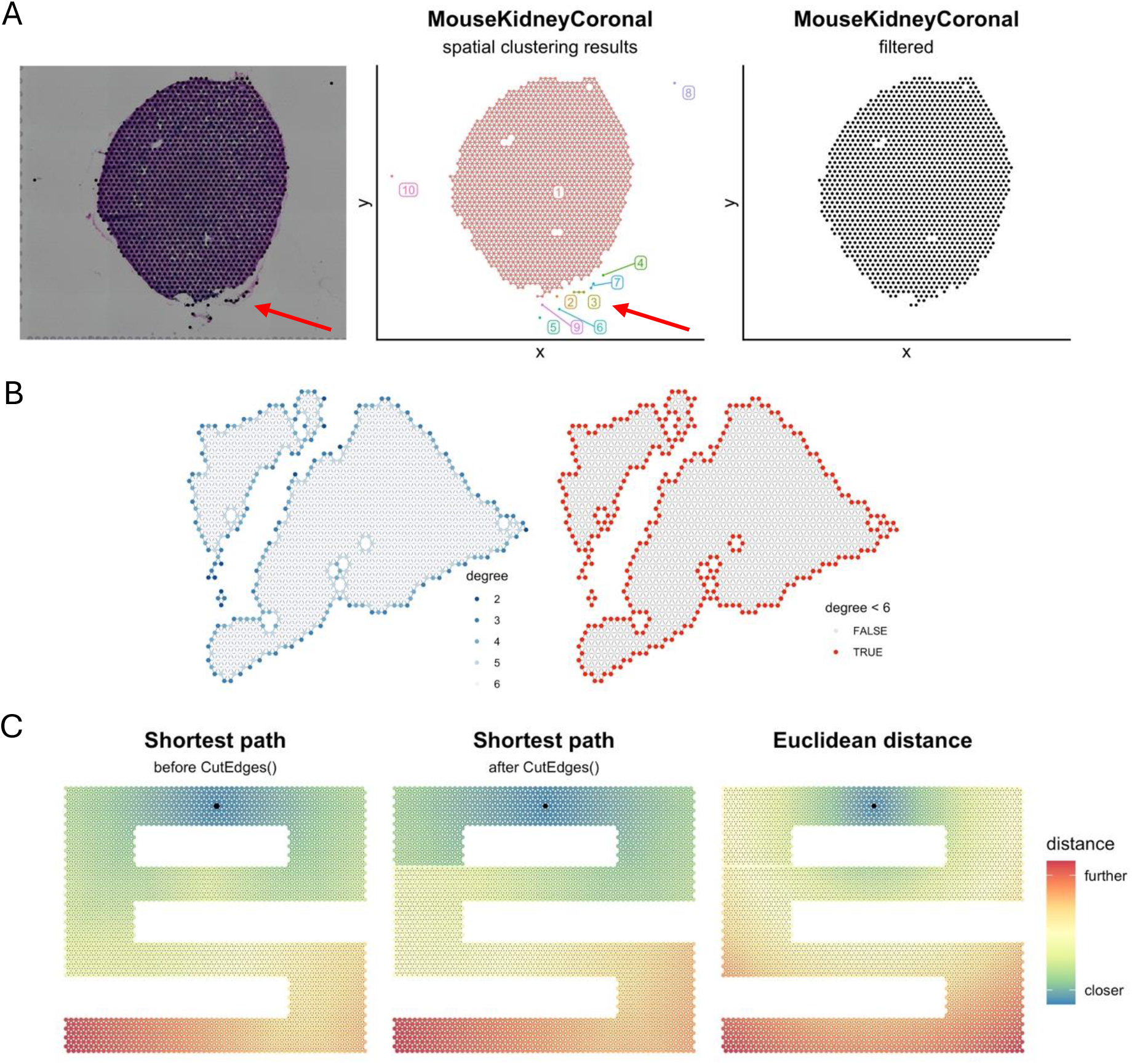
Useful scenarios for SpotGraphs. (A) H&E staining with corresponding spots from the TENxVisium MouseKidneyCoronal dataset. Spatial clustering in the resulting igraph object identifies groups of connected spots based on adjacency, which can be used for spot-level filtering. Red arrows point to low quality spots (B) The degree (i.e., number of connections) of each spot can identify which spots are at the boundaries of the tissue sample. (C) Distance from black spot to every other spot calculated by shortest path before (left panel) and after removing some connections (middle panel) with CutEdges(), or by Euclidean distance (right panel).

### Identifying boundary spots

With igraph, we can identify the spots laying at the boundaries of our tissue sample by calculating the degree of each spot. In a grid, any spot with a degree less than the maximum degree of the entire graph will be at some tissue boundary. In other words, spots at the boundaries of our tissue should have fewer adjacent spots than a spot at the center of the tissue sample that is fully surrounded at the center of a hexagonal or square grid. For example, the maximum degree for a hexagonal grid would be 6 adjacent spots, so any spot with 5 or fewer adjacent spots can be considered to be at the boundary (Figure 3B).

### Calculating distance via shortest path

Alternative distance metrics can be calculated once an igraph object is constructed, for example, if the tissue sample is distorted in a way where Euclidean distance does not provide accurate estimates (Figure 3C - right panel). In these cases, the analyst can create an igraph object with SpotGraph() to calculate the shortest path along the tissue using the igraph::distances() function (Figure 3C - middle panel). A user can also modify the connections between spots with the CutEdges() function (Figure 3C - left panel), which will remove all edges connecting two groups of spots to adjust the distances calculated with igraph::distances().

## Discussion

Through the SpotGraphs R package, we provide a set of tools to directly interact with the x,y-coordinates of spatial transcriptomics data in a more flexible manner by interfacing with the existing igraph infrastructure in R. We provide examples where easy access to igraph functions, enabled by SpotGraphs, allows users to fine-tune their spot-level filtering, identify neighborhood centers and boundaries, and calculate alternative distance metrics, but we anticipate many other scenarios where access to igraph through our package will further enhance spatial analysis for users in R. We also find that in situations where the number of adjacent neighbors or the overall structure of the adjacency network can influence the results of downstream analyses, the nearest-neighbor approach should be avoided entirely. For example, if a user would like to calculate the distance between two spots using the shortest path through the adjacency network, spots at the boundaries would artificially seem closer to each other than spots further away from the tissue boundaries. Instead, we recommend using either our Euclidean distance or Delaunay triangulation based approaches to build the adjacency matrices in R, or the SquidPy graph API in Python. A limitation of this study is that we have not benchmarked ST data from platforms that generate data that is not organized in a grid, such as 10X Xenium, Slide-Seq, and MERFISH, but in these data types, Delaunay triangulation has been shown to perform reasonably well (Dries et al., 2021; Miller et al., 2021).

## References

Antonov, M., Csárdi, G., Horvát, S., Müller, K., Nepusz, T., Noom, D., Salmon, M., Traag, V., Welles, B. F., & Zanini, F. (2023). Igraph enables fast and robust network analysis across programming languages (arXiv:2311.10260). arXiv. 10.48550/arXiv.2311.10260

Arora, R., Cao, C., Kumar, M., Sinha, S., Chanda, A., McNeil, R., Samuel, D., Arora, R.K., Matthews, T. W., Chandarana, S., Hart, R., Dort, J. C., Biernaskie, J., Neri, P., Hyrcza, M. D., & Bose, P. (2023). Spatial transcriptomics reveals distinct and conserved tumor core and edge architectures that predict survival and targeted therapy response. Nature Communications, 14(1), 5029. 10.1038/s41467-023-40271-4

Crowell, H. (2025). TENxVisiumData: Visium spatial gene expression data by 10X Genomics. Bioconductor. http://bioconductor.org/packages/TENxVisiumData/

Csardi, G., & Nepusz, T. (2006). The igraph software package for complex network research. InterJournal, Complex Systems.

Csárdi, G., Nepusz, T., Müller, K., Horvát, S., Traag, V., Zanini, F., & Noom, D. (2025). igraph for R: R interface of the igraph library for graph theory and network analysis [Computer software]. Zenodo. 10.5281/zenodo.17470081

Dries, R., Zhu, Q., Dong, R., Eng, C.-H. L., Li, H., Liu, K., Fu, Y., Zhao, T., Sarkar, A., Bao, F., George, R. E., Pierson, N., Cai, L., & Yuan, G.-C. (2021). Giotto: A toolbox for integrative analysis and visualization of spatial expression data. Genome Biology, 22(1), 78. 10.1186/s13059-021-02286-2

Gebhardt, A., Bivand, R., & library), D. S. (author of the shull. (2024). interp: Interpolation Methods (Version 1. 1-6) [Computer software]. https://cran.r-project.org/web/packages/interp/index.html

Jefferis, G., Kemp, S. E., Müller, K., Arya [aut, S.s s, cph, Mount [aut s, D., cph, & details),U. of M. (ANN library is copyright U.s of M. and S. A. and D. M. S. file C. for. (2024).

RANN: Fast Nearest Neighbour Search (Wraps ANN Library) Using L2 Metric (Version 2.6.2) [Computer software]. https://cran.r-project.org/web/packages/RANN/index.html

Kalinowski, T., Ushey, K., Allaire, J. J., RStudio, Tang [aut, Y., cph, Eddelbuettel, D., Lewis, B., Keydana, S., Hafen, R., library, M. G. (TinyThread, & http://tinythreadpp.bitsnbites.eu/). (2025). reticulate: Interface to “Python” (Version 1.44.1) [Computer software]. https://cran.r-project.org/web/packages/reticulate/index.html

Miller, B. F., Bambah-Mukku, D., Dulac, C., Zhuang, X., & Fan, J. (2021). Characterizing spatial gene expression heterogeneity in spatially resolved single-cell transcriptomic data with nonuniform cellular densities. Genome Research, 31(10), 1843–1855. 10.1101/gr.271288.120

Palla, G., Spitzer, H., Klein, M., Fischer, D., Schaar, A. C., Kuemmerle, L. B., Rybakov, S., Ibarra, I. L., Holmberg, O., Virshup, I., Lotfollahi, M., Richter, S., & Theis, F. J. (2022). Squidpy: A scalable framework for spatial omics analysis. Nature Methods, 19(2), 171–178. 10.1038/s41592-021-01358-2

Salim, A., Bhuva, D. D., Chen, C., Yang, P., Davis, M. J., & Yang, J. Y. (2024). SpaNorm: Spatially-aware normalisation for spatial transcriptomics data. 10.1101/2024.05.31.596908

Yan, G., Hua, S. H., & Li, J. J. (2025). Categorization of 34 computational methods to detect spatially variable genes from spatially resolved transcriptomics data. Nature Communications, 16(1), 1141. 10.1038/s41467-025-56080-w

Zhao, E., Stone, M. R., Ren, X., Guenthoer, J., Smythe, K. S., Pulliam, T., Williams, S.R., Uytingco, C. R., Taylor, S. E. B., Nghiem, P., Bielas, J. H., & Gottardo, R. (2021).Spatial transcriptomics at subspot resolution with BayesSpace. Nature Biotechnology, 39(11), 1375–1384. 10.1038/s41587-021-00935-2

